# Sex-specific effects of polygenic risk for schizophrenia on lifespan cognitive functioning in healthy individuals

**DOI:** 10.1101/2021.06.23.449554

**Authors:** Elise Koch, Lars Nyberg, Anders Lundquist, Sara Pudas, Rolf Adolfsson, Karolina Kauppi

## Abstract

Polygenic risk for schizophrenia has been associated with lower cognitive ability and age-related cognitive change in healthy individuals. Despite well-established neuropsychological sex differences in schizophrenia patients, genetic studies on sex differences in schizophrenia in relation to cognitive phenotypes are scarce. Here, we investigated whether the effect of a polygenic risk score (PRS) for schizophrenia on childhood, midlife and late life cognitive function in healthy individuals is modified by sex, and if PRS is linked to accelerated cognitive decline. Using a longitudinal data set from healthy individuals aged 25-100 years (N = 1,459) spanning a 25-year period, we found that PRS was associated with lower cognitive ability (episodic memory, semantic memory, visuospatial ability), but not with accelerated cognitive decline. A significant interaction effect between sex and PRS was seen on cognitive task performance, and sex-stratified analyses showed that the effect of PRS was male-specific. In a sub-sample, we observed a male-specific effect of the PRS on school performance at age 12 (N = 496). Our findings of sex-specific effects of schizophrenia genetics on cognitive functioning across the life-span indicate that the effects of underlying disease genetics on cognitive functioning is dependent on biological processes that differ between the sexes.

## Introduction

Schizophrenia is a severe neuropsychiatric disorder that affects about 1% of the population^1,2^. Cognitive deficits are considered a core feature of schizophrenia, affecting about 80% of the patients^1,3^. The cognitive symptoms in schizophrenia involve working memory, episodic memory, reasoning and problem solving, speed of processing, and social cognition^3,4^. Cognitive deficits in schizophrenia have been observed before the onset of positive symptoms, indicating that cognitive symptoms are not only related to secondary effects of the disease such as antipsychotic medication^5,6^. Schizophrenia has also been suggested to be associated with accelerated cognitive aging^7^. To what extent this link may be due to disease-related factors, such as psychosis or taking antipsychotics, or underlying genetics is poorly understood. Evidence from studies reporting a link between poor school performance and risk of developing schizophrenia later in life^8,9^ support the possibility that the cognitive deficits in schizophrenia patients may arise due to developmental causes already at young age^10^. However, it is not known if schizophrenia genetics is related to school performance in the general population.

Sex differences have long been observed in schizophrenia patients, with males being affected more frequently and more severely than females^11,12^. Moreover, male patients have an earlier age at onset, worse negative symptoms, and worse treatment response to antipsychotics compared to female patients^13–16^. Several studies reported that male patients have more severe cognitive impairment than female patients^17–22^, but other studies have found the opposite^23,24^ or no sex differences in cognitive deficits in schizophrenia^25^. Proposed explanations for sex differences in cognitive abilities in schizophrenia include sex hormone differences^26,27^ and sex differences in brain structure and volume^27,28^.

Genetics is known to play a crucial role in the development of schizophrenia, with an estimated heritability of approximately 80%^1,2^. The schizophrenia genome-wide association study (GWAS) performed by the Psychiatric Genomics Consortium (PGC)^29^ has identified a large number of genetic risk variants, typically single nucleotide polymorphisms (SNPs), that individually have weak effects on the phenotype^30^. To capture the polygenic nature of schizophrenia, a polygenic risk score (PRS) can be calculated to examine the impact of cumulative genetic risk for schizophrenia on related phenotypes^31,32^. Studying PRS in unrelated healthy individuals has the advantage that genetic effects can be separated from secondary disease-related factors such as medication. A genetic overlap between schizophrenia and cognitive functioning in healthy individuals has been identified^33–35^ and schizophrenia PRS have been linked to decreased cognitive ability across different cognitive domains in both schizophrenia patients^34,36^ and healthy individuals^37,38^. However, sex differences in relation to cognitive phenotypes were typically not reported in past genetic studies of schizophrenia^34,36–39^, but a recent study suggests male-specific effects of schizophrenia PRS on memory in healthy older adults^40^.

In the present study, we investigated sex-specific effects of schizophrenia PRS on cognitive performance in healthy individuals. We used longitudinal cognitive data from a large sample of healthy adults (25-100 years) to investigate genetic effects on both cognitive level and slope (episodic memory, semantic memory, visuospatial ability). In a sub-sample, we examined the impact of PRS on school grade data at age 12 as a proxy for childhood cognition.

## Methods

### Participants

Data in the present study come from the longitudinal population-based Betula Prospective Cohort Study on memory, health and aging, conducted in Umeå, Sweden^41,42^, including measurements of cognitive functions from six test waves (T1-T6) five years apart, with a total follow-up period of 25 years. Exclusion criteria were dementia and known neurologic or psychiatric disease. From three cohorts (S1, S3 and S6), a total of 1,746 individuals had been successfully genotyped. We excluded individuals that had developed dementia (N = 287), resulting in 1,459 individuals (with an average of 3.6 time-points, ranging from 1-6) included in the current study (678 males, 781 females) aged between 25-100 years (see **Supplementary Table 1** for age distribution by time from inclusion). Dementia diagnosis was done by a geropsychiatrist based on the DSM-IV criteria^43^ as previously described elsewhere^41,44,45^. All participants had European ancestry. The research was approved by the regional ethical review board at Umeå University (EPN), and all participants gave written informed consent.

**Table 1:**
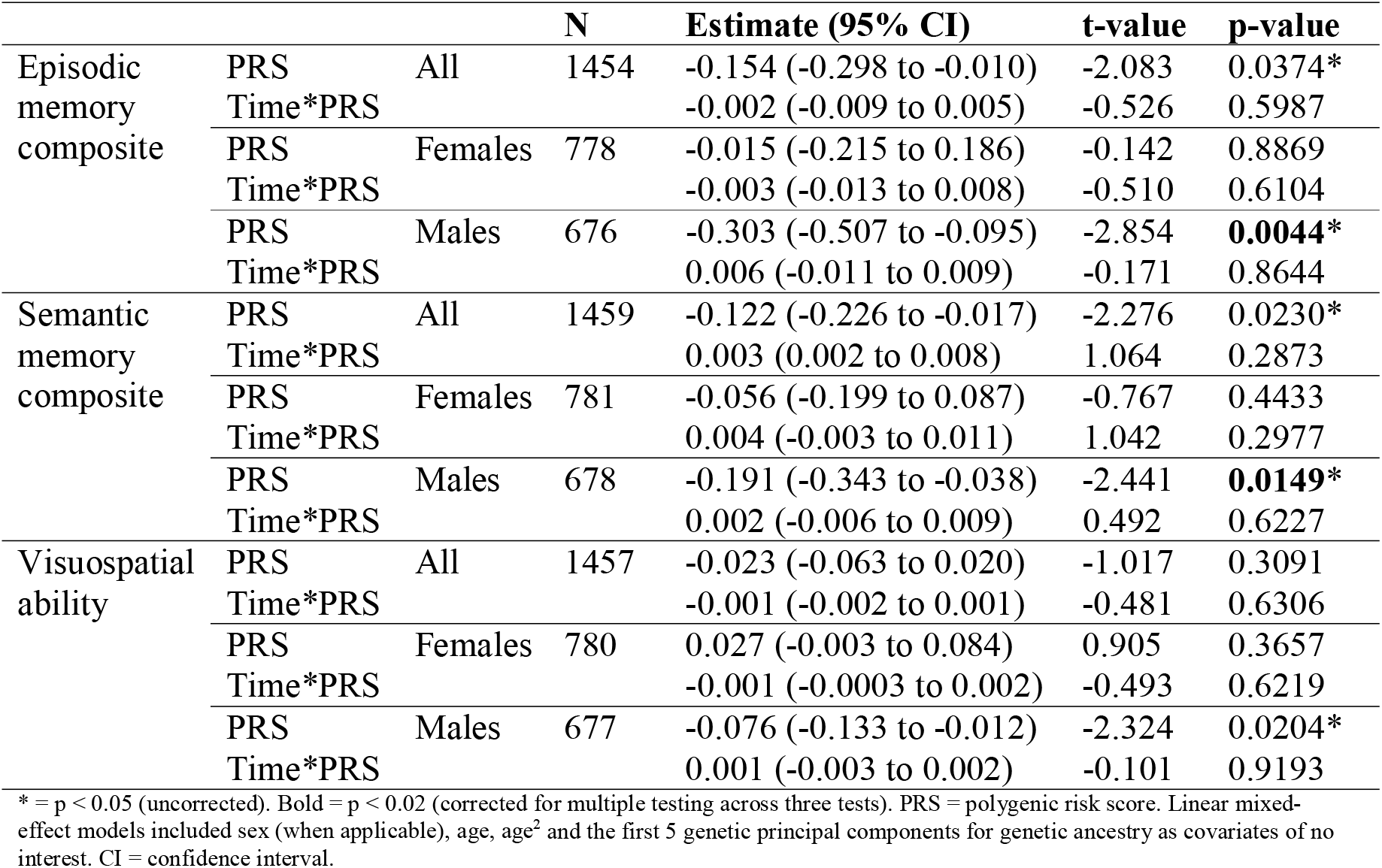
Effect of schizophrenia PRS (p-value threshold ≤ 1) on intercept and slope of individual cognitive tests, including all six test occasions (T1-T6).

### Genotyping and construction of polygenic risk scores

Two genotyping datasets were used in the present study. For the first dataset, most of the DNA extraction for SNP genotyping was done at VIB-U Antwerp Center for Molecular Neurology in Belgium, and a minor part of DNA samples was extracted at Genome-wide genotyping LGC Genomics Ltd, UK. All DNA samples were genotyped using two types of Illumina arrays: The Infinium Exome Array and the Infinium Human OmniExpress-12v1_H Array. This was done at the Genotyping Platform of the Broad Institute of MIT and Harvard, USA between 2012 – 2014. Imputation and standard quality control (QC) of the raw genotypes was done using the 1000 Genomes data and the imputation pipeline RICOPILI^46^ used by the psychiatric genomic consortia (PGC). For the second genotyping dataset, the DNA extraction for SNP genotyping was done at the Institute of Human Genetics, University of Bonn, Germany. All DNA samples were genotyped using two types of Illumina arrays: Illumina Omni Express and Omni 1S Bead chip kits. Imputation of the raw genotypes was done according to the ENIGMA2 protocol of the ENIGMA Consortium (http://enigma.ini.usc.edu/) to the 1000 genomes reference panel^47^ using minimac tools^48^. For both genotyping datasets, post-imputation QC was performed based on genotype call rate < 10%, minor allele frequency (MAF) < 1%, SNP missingness < 5 %, and imputation info < 0.8. After removal of SNPs with ambiguous strand alignment and strand mismatch between the two datasets, we merged these datasets using PLINK^49^ (version 1.9). Thereafter, schizophrenia PRS were calculated in PLINK^49^ based on the summary statistics of the PGC2 schizophrenia GWAS^29^ including 35,476 cases and 46,839 controls, after excluding the current sample as well as SNPs within the extended major histocompatibility complex (MHC) region (25-34 Mb on chromosome 6 on the hg19 assembly) due to the high linkage disequilibrium (LD) of this region. First, LD clumping was performed according to parameters used by the PGC2^29^: discarding SNPs within 500 kb of, and in r^2^ ≥ 0.1 with another more significant SNP, and excluding SNPs with MAF < 10%, after removal of SNPs with genotype call rate < 1% and MAF < 1%. PRS were based on SNPs located on autosomal chromosomes and calculated for each individual by summing the alleles of the clumped SNPs (N = 75,992) weighted by the natural log of the odds ratio from the PGC2 GWAS results^29^. The explained variance in case-control discrimination has been shown to increase with increasing PRS p-value threshold, reaching a plateau at p < 0.05^29,50^. However, the p-value threshold explaining the largest variance in case-control discrimination may not be the same as that for the amount of variance explained in endophenotypes. Moreover, it has been shown that the p-value threshold of 1 has the highest empirical power in traits with high polygenicity^51^, and both schizophrenia^2^ and cognitive performance^52^ show highly polygenic inheritance. Therefore, PRS were constructed at 10 p-value thresholds (0.0001, 0.001, 0.01, 0.05, 0.1, 0.2, 0.3, 0.4, 0.5, 1), and the p-value threshold that explained the largest amount of variance in the cognitive composite cross-sectionally, using Nagelkerke’s pseudo-R^2^, was used in our main analyses. Additional PRS p-value thresholds were reported for comparisons and transparency, as the optimal p-value threshold may differ depending on the outcome variable. In addition, we calculated a polygenic score (PGS) on cognitive performance in the same way as the schizophrenia PRS using summary statistics from a large multicenter GWAS on cognitive performance measured across at least three cognitive domains including 257,841 individuals^52^, and we used the PGS including all clumped SNPs (N = 75,639) from the GWAS summary statistics (p-value threshold ≤ 1).

### Cognitive tests

The analyses were based on a cognition composite score that was calculated as the sum of z-transformed tests of visuospatial ability, episodic memory and semantic memory, as well as on the three cognitive domains separately. For episodic memory, a composite score was calculated as the sum of z-transformed tests of episodic memory (two tests of free oral recall of verb-noun sentences, two tests of category-cued recall of nouns from the sentence recall, and one test of free recall of presented nouns). For semantic memory, a composite score was calculated as the sum of z-transformed tests of semantic knowledge/verbal fluency (three tests of verbal generation of as many words as possible during 1 min: words that begin with A; Words that begin with M, exactly 5 letters; professions that begin with B). Visuospatial ability was measured with the block design task from the Wechsler Adult Intelligence Scale (WAIS-R)^53^. For details about cognitive tasks see Nilsson et al. 2004^41^.

### School performance

School grade data at approximately age 12 (sixth grade) was available for a subset of 496 Betula participants (247 males and 249 females) from the cohorts S1 and S3. The analyses were based on a school grade composite score that was calculated as the sum of z-transformed school grades from six school subjects: mathematics, Swedish, history, biology, geography, and scripture knowledge (Christianity) as well as on the six grades separately. For details see Pudas et al. 2019^54^.

### Statistical analyses

To examine the association of PRS with level and change in cognitive measures over 25 years, we performed linear mixed-effect models that were fitted in R using the lme4 function available through the lme4 package. P-values were estimated based on the Satterthwaite approximations implemented in the lme4Test package. The models included the PRS as covariate of interest and the following covariates of no interest: age, age^2^, sex, and the first 5 principal components for genetic ancestry (to control for population stratification). Time from inclusion (years) was used as time-scale to represent slope, and interaction with time was allowed for all covariates. The models also included random subject-specific intercepts. In addition, we ran the same models adding a polygenic score (PGS) for cognitive performance. Model comparisons were done with a Likelihood-Ratio-Test. To test for interaction effects, an interaction term for age and PRS (PRS*age) as well as for sex and PRS (PRS*sex) was included in the longitudinal regression models. Cross-sectional analyses of school performance at age 12 were done using the lm-function in R including the PRS as well as birth month (to account for differences in cognitive maturity), sex, and the first 5 principal components. Both the longitudinal and the cross-sectional analyses were performed for all individuals as well as sex-stratified. The variance in cognitive task performance explained by PRS was calculated as Nagelkerke’s pseudo-R^2^, by comparing the full model (PRS plus covariates) to the reduced model (covariates only), which was done for cross-sectional models using the lm-function in R. These models were based on each individual’s first cognitive test occasion, and included sample (cohort) as additional covariate. Levene’s test was used to assess the equality of variance in baseline cognitive test performance and schizophrenia PRS in males and females. Welch Two Sample t-tests were performed to investigate if males and females differ according to their cognitive test performance at baseline as well as their PRS. A logistic regression analysis was performed to test if intake of antipsychotics is associated with PRS. All statistical analyses were performed in R version 4.0.3.

## Results

Descriptive statistics for each cognitive test and test occasion (T1-T6) are shown in **Supplementary Table 2** separately for males and females as well as for all individuals. The effect of schizophrenia PRS (calculated with 10 different p-value thresholds) on cognition composite at baseline can be found in **Supplementary Table 3**, showing that the variance in cognitive test performance explained by schizophrenia PRS is largest for the PRS calculated with a p-value threshold ≤ 1, which was used for subsequent analyses. Using the cognition composite, Levene’s test showed that males and females had equal variance in their cognitive test performance (f = 2.659, p-value = 0.103). Consistent with reported cognitive sexual dimorphisms^55^, Welch Two Sample t-tests showed that males performed better on the visuospatial task (t = −3.7691, df = 1423.3, p-value = 0.0002), whereas females performed better on the episodic memory (t = 4.3799, df = 1443.7, p-value = 1.273e−05) and semantic memory tasks (t = 4.7805, df = 1427.9, p-value = 1.929e−06), shown in **Figure 1**. Moreover, Welch Two Sample t-test showed that males and females did not differ significantly regarding their schizophrenia PRS (t = 1.888, df = 1419.8, p-value = 0.059, males having a slightly higher PRS). Levene’s test showed that males and females had equal variance in their PRS (f = 0.012, p-value = 0.9173).

**Table 2:** Effect of schizophrenia PRS (p-value threshold ≤ 1) on each school grade at age 12, separately for males and females as well as for all individuals.

|  |  | N | Estimate (95% CI) | t-value | p-value |
| --- | --- | --- | --- | --- | --- |
| Mathematics | All | 496 | -0.082 (-0.170 to 0.006) | -1.837 | 0.0668 |
|  | Females | 249 | -0.012 (-0.132 to 0.1081) | -0.195 | 0.8457 |
|  | Males | 247 | -0.170 (-0.299 to -0.041) | -2.603 | 0.0098* |
| Swedish | All | 496 | -0.033 (-0.119 to 0.054) | -0.748 | 0.4549 |
|  | Females | 249 | 0.030 (-0.091 to 0.152) | 0.491 | 0.6240 |
|  | Males | 247 | -0.111 (-0.234 to 0.013) | -1.767 | 0.0785 |
| History | All | 496 | -0.100 (-0.189 to -0.011) | -2.208 | 0.0278* |
|  | Females | 249 | -0.018 (-0.142 to 0.106) | -0.281 | 0.7788 |
|  | Males | 247 | -0.195 (-0.324 to -0.066) | -2.969 | **0.0033\*** |
| Biology | All | 496 | -0.077 (-0.167 to 0.013) | -1.683 | 0.0931 |
|  | Females | 249 | 0.042 (-0.081 to 0.165) | 0.676 | 0.4999 |
|  | Males | 247 | -0.214 (-0.344 to -0.084) | -3.236 | **0.0014\*** |
| Geography | All | 496 | -0.050 (-0.139 to 0.039) | -1.110 | 0.2678 |
|  | Females | 249 | 0.081 (-0.039 to 0.202) | 1.331 | 0.1845 |
|  | Males | 247 | -0.202 (-0.332 to -0.072) | -3.061 | **0.0025\*** |
| Scripture knowledge (Christianity) | All | 496 | -0.096 (-0.181 to -0.012) | -2.239 | 0.0256* |
|  | Females | 249 | -0.028 (-0.145 to 0.089) | -0.474 | 0.6357 |
|  | Males | 247 | -0.186 (-0.308 to -0.064) | -3.005 | **0.0029\*** |

\* = $p < 0.05$ (uncorrected). Bold = $p < 0.008$ (corrected for multiple testing across six subjects). PRS = polygenic risk score. Linear regression models included sex (when applicable), birth month and the first 5 genetic principal components for genetic ancestry as covariates of no interest. CI = confidence interval.

**Figure 1:**
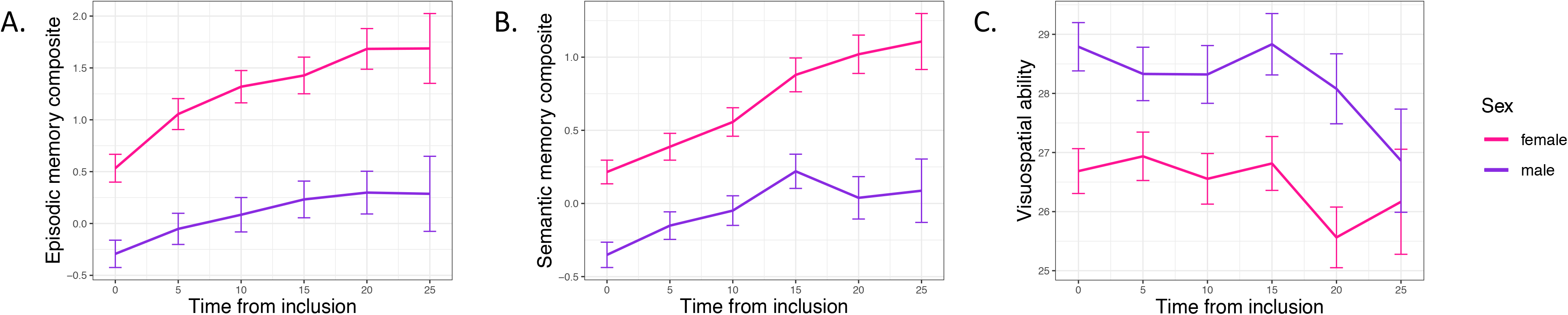
Cognitive test performance for each cognitive domain across 25 years separately for males and females. A. Episodic memory. B. Semantic memory. C. Visuospatial ability. Error bars show standard error.

### Effect of schizophrenia genetics on cognitive test performance

Overall, schizophrenia PRS was associated with a lower cognition composite (t-value = − 2.224, p-value = 0.026), but not with cognitive change over time (t-value = 0.188, p-value = 0.851). Results subdivided by three cognitive domains showed that PRS was negatively associated with episodic memory and semantic memory, but not with visuospatial ability (**Table 1**). For any cognitive domain, no significant effects of PRS on cognitive change over time could be observed (**Table 1**). Complementary analyses of PRS using more stringent p-value thresholds again showed no effects on visuospatial ability, nor on cognitive slope in any examined domain (**Supplementary Table 4**).

Next, we tested for interaction effects of PRS with age or sex on cognitive test performance cross-sectionally using the cognition composite within linear models. There was no significant interaction between PRS and age (t-value = 1.789, p-value = 0.074), but a significant interaction effect between PRS and sex could be observed (t-value = −2.262, p-value = 0.024). There was a significant interaction effect between PRS and sex for all 10 PRS calculated with different p-value thresholds for the cognition composite (**Supplementary Table 5**). Sex-stratified analyses showed that the association between PRS and lower cognitive performance across all cognitive domains was male-specific with no trend seen in females (**Table 1, Figure 2A**) and again, no longitudinal effect of PRS on cognitive performance could be observed in either sex (**Table 1).** As a sensitivity analysis, we also included a polygenic score for cognitive performance (Cog-PGS) in the models to see if the effects of schizophrenia genetics are independent of genetic variants with a known effect on cognition, based on GWAS on cognitive ability in healthy individuals^52^. As expected, the Cog-PGS was strongly predictive of cognitive ability (**Supplementary Table 6**), but no interaction effects with sex were seen on the cognition composite score (t-value = −0.628, p-value = 0.530). Including all individuals, the negative association between the schizophrenia PRS and cognitive performance was no longer significant when also including the Cog-PGS in the same model. Model comparisons showed that models including the Cog-PGS alone did not differ significantly regarding fit to data from the models including both the Cog-PGS and schizophrenia PRS (Episodic memory: χ^2^ = 3.888 (2), p = 0.143; Semantic memory: χ^2^ = 3.272 (2), p = 0.195; Visuospatial ability: χ^2^ = 0.643 (2), p = 0.725). However, sex-stratified analyses showed that the negative association between schizophrenia PRS and cognitive performance (episodic memory and semantic memory) was still significant in males when including the Cog-PGS in the models (**Supplementary Table 6**). In another sensitivity analysis, we ran the models including years of education as additional covariate showing that education was strongly predictive for cognitive performance, but the relationship between schizophrenia PRS and cognitive performance was independent of participants’ education (shown for all individuals as well as sex-stratified in **Supplementary Table 7**). Data on self-reported drug use was available for T3-T6. Logistic regression analysis showed that intake of antipsychotics (N = 14) was associated with higher PRS (OR = 3.204, p-value = 3.65e−06). To test if the effect of schizophrenia genetics on cognitive performance is independent of antipsychotic use, we ran the models excluding individuals taking one or more antipsychotics (N = 14). For all three cognitive domains, the significance level of the PRS as well as the interaction effect with sex did not change.

**Figure 2:**
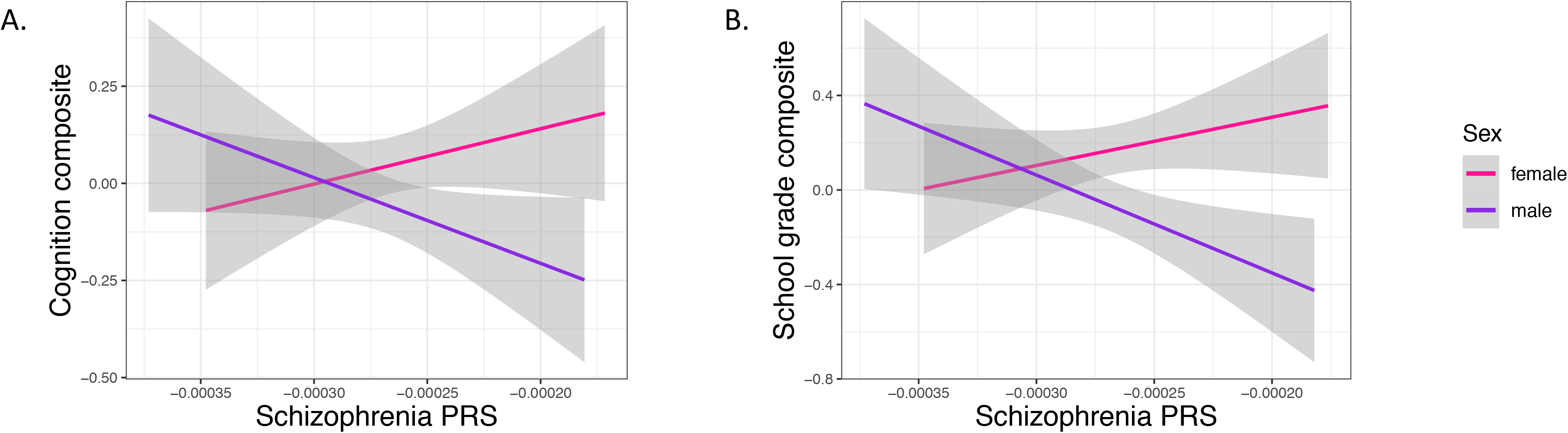
Association between schizophrenia PRS and cognitive test performance in males and females. **A.** PRS associated with a cognition composite score that was calculated as the sum of z-transformed tests of visuospatial ability, episodic memory, and semantic memory. **B.** PRS associated with a school grade composite score that was calculated as the sum of z-transformed school grades from six school subjects: mathematics, Swedish, history, biology, geography, and scripture knowledge (Christianity). Grey areas indicate standard error. PRS p-value threshold ≤ 1.

### Effect of schizophrenia genetics on school performance

We tested for association between schizophrenia PRS and school performance at age 12 (N = 496), including sex interactions, with school grades used as a proxy for childhood cognitive ability. There was a trend for an association between PRS and lower school grade composite across six school subjects (t-value = −1.908, p-value = 0.0570). Importantly, there was a significant interaction between PRS and sex (t-value = −2.962, p-value = 0.0032). Sex-stratified analyses showed the same patterns as for cognition in mid- and old age; a significant negative association in males (t-value = −3.235, p-value = 0.0014), with no trend in females (t-value = 0.305, p-value = 0.7604) (**Figure 2B**). Results subdivided by each school subject separately showed the same patterns with no trend for females in any school subject (**Table 2**).

## Discussion

Using a large sample of healthy individuals, we found a robust interaction effect between schizophrenia PRS and sex on cognitive ability. A male-specific effect of the PRS was seen on school grades at age 12 across 6 subjects and on cognitive test performance across three domains, episodic memory, semantic memory, visuospatial ability in mid- to old age. Using a longitudinal data set spanning a 25-year period, we also showed that PRS was not associated with accelerated cognitive decline.

Sex differences can be described as sex-specific effects (presence in one sex only) or sex-dependent (quantitative differences between the sexes)^56^. In the present study, the association between the PRS and worse cognitive performance in mid- and old age, as well as worse school performance was seen in males only with no trend effects in females, indicating sex-specific effects of schizophrenia genetics on cognitive ability in healthy individuals across the entire lifespan. Although it has been proposed that the impact of sex should be tested in genetic studies of schizophrenia given the significant differences in disease manifestation^56–58^, to our knowledge, sex differences in the effect of schizophrenia PRS on cognition has only been considered in one previous study, where sex-stratified analyses showed that the effect of schizophrenia PRS on verbal memory and semantic fluency in older adults reached significance only in males^40^. Here, we were able to demonstrate a significant sex*PRS interaction thus demonstrating that there are statistically reliable differences in the effect of PRS between the sexes, and showed that this effect was not seen for a genetic score for general cognitive ability, but instead specific to schizophrenia genetics. In line with reported cognitive sexual dimorphisms^55^, we show that males performed better on the visuospatial task, whereas females performed better on the memory tasks. Thus, our observations of a male-specific effect of schizophrenia genetics on both memory tasks and visuospatial ability indicate that this effect is independent of normal cognitive sexual dimorphisms. Finally, we show that male-specificity not only relates to cognitive functioning in adulthood and old age, but also to childhood cognitive ability.

A schizophrenia GWAS stratified by sex did not reveal any genome-wide significant sex-specific associations^59^, but a genome-wide genotype-by-sex interaction analysis of risk for schizophrenia found genome-wide significant SNP-by-sex interactions^57^. However, no consistent sex differences in SNP-based heritability have been identified in schizophrenia^58^. Taken together, our results suggest that the effect of schizophrenia genetics on a certain disease endophenotype, cognitive ability, is sex-specific. As the effect of schizophrenia PRS on cognition might differ between healthy individuals and schizophrenia patients^60^, our results warrant replication in patient cohorts to examine if sex differences are related to underlying disease genetics on specific symptoms and disease manifestations. Further, it will be relevant to not only replicate the observed sex-specific PRS effect for episodic memory, semantic memory, and visuospatial ability as investigated here, but also to explore its generality in other cognitive domains. Finally, to investigate if the male-specific effects on cognition are driven by specific biological pathways of schizophrenia, future studies investigating pathway-specific PRS would be of importance.

Using a long follow-up time, we did not observe any association between schizophrenia PRS and accelerated cognitive decline. These results are in contrast to a previous publication by McIntosh et al.^61^ where schizophrenia PRS was associated with change in cognitive ability from childhood to old age in healthy individuals^61^. In a previous study of older adults, only a modest decline in overall cognitive performance associated with schizophrenia PRS could be observed^62^, and in another two longitudinal studies of older adults^40,63^, no such association was found. In our sample, there was a trend for a negative interaction between age and PRS on cognition (p = 0.07), indicating that the effect of PRS on cognitive task performance may be somewhat stronger in young individuals. The cognitive deficits in schizophrenia patients may arise at younger age due to developmental causes, as also our school grade data suggest. Indeed, it has been shown that children who later developed schizophrenia had deficits in tests of processing speed, attention, visuospatial ability, and working memory when they entered school at age 7, and were further impaired in attention and working memory as they got older compared to children that did not develop schizophrenia^64^. This was further confirmed by our finding of worse school performance at age 12 in relation to schizophrenia PRS. However, it has also been reported that schizophrenia PRS has no effect on the association between low school performance and developing schizophrenia^65^, which could be due to sex differences that were not considered in that study.

Pharmacological treatment options mainly include antipsychotics that primarily treat the positive symptoms^66^, and besides non-pharmacological treatment options there are currently no available medications that efficiently treat the cognitive symptoms in schizophrenia^67^. As sex differences observed in cognitive performance in schizophrenia may be due to differences in sex hormone levels, estradiol and other sex hormones have been suggested for the treatment of cognitive symptoms in schizophrenia^68,69^. The selective estrogen receptor modulator raloxifene in combination with antipsychotics has shown beneficial effects on cognition in both women and men with schizophrenia^68–70^. However, the genetic mechanisms of sex differences related to cognitive processing in schizophrenia require further investigation. It should be tested if our results translate to schizophrenia patients, which may enable the development of sex-specific pharmocological treatments for schizophrenia and a greater understanding of the biology of cognitive impairments in the disease.

In conclusion, our results strongly suggest male-specific effects of schizophrenia genetics on episodic memory, semantic memory, and visuospatial ability in healthy individuals. Our findings indicate that the effect of underlying schizophrenia genetics on cognition is dependent on biological processes that differ between the sexes. These results open up for replication in patients and further investigations of interactions between schizophrenia genetics and biological processes with known sex differences, which could provide valuable insight into disease biology, and potentially lead to the development of novel sex-specific pharmacological treatment options.

## Funding

This work was supported by a grant to KK from the Swedish Research Council (Grant no 2017-03011). The retrieval of school grades was supported by a grant from the Royal Swedish Academy of Sciences (AS2015-0004) to SP.

## Conflicts of interest

The authors have no conflicts of interest to declare.

## Notes

### Competing Interest Statement

The authors have declared no competing interest.

