## Supplementary Materials for "Sex-specific effects of polygenic risk for schizophrenia on lifespan cognitive functioning in healthy individuals"

^1^Department of Integrative Medical Biology, Umeå University, Sweden, ^2^Department of Radiation Sciences, Diagnostic Radiology, University Hospital, Umeå University, Sweden, ^3^Department of Statistics, School of Business, Economics and Statistics, Umeå University, Sweden, ^4^Department of Clinical Sciences, Umeå University, Sweden, ^5^Department of Medical Epidemiology and Biostatistics, Karolinska Institute, Sweden

**Supplementary Table 1.** Age distribution of participants by time from inclusion (years). Drop-out by age-bins can be read diagonally, as there were five years between both time points and age bins.

|  | **AGE** |  |  |  |  |  |  |  |  |  |  |  |  |  |  |  |  |
| --- | --- | --- | --- | --- | --- | --- | --- | --- | --- | --- | --- | --- | --- | --- | --- | --- | --- |
| **TIME** | **25** | **30** | **35** | **40** | **45** | **50** | **55** | **60** | **65** | **70** | **75** | **80** | **85** | **90** | **95** | **100** | **sum** |
| **0** | 10 | 9 | 96 | 188 | 185 | 185 | 163 | 158 | 119 | 102 | 109 | 92 | 43 | 0 | 0 | 0 | **1459** |
| **5** | 0 | 6 | 7 | 92 | 177 | 166 | 170 | 153 | 145 | 109 | 93 | 92 | 72 | 20 | 0 | 0 | **1302** |
| **10** | 0 | 0 | 0 | 0 | 77 | 153 | 144 | 142 | 136 | 133 | 86 | 65 | 44 | 42 | 5 | 0 | **1027** |
| **15** | 0 | 0 | 0 | 0 | 0 | 67 | 130 | 120 | 133 | 120 | 98 | 55 | 41 | 25 | 15 | 0 | **804** |
| **20** | 0 | 0 | 0 | 0 | 0 | 0 | 59 | 103 | 98 | 108 | 90 | 66 | 23 | 15 | 7 | 3 | **572** |
| **25** | 0 | 0 | 0 | 0 | 0 | 0 | 0 | 43 | 39 | 43 | 43 | 31 | 15 | 3 | 2 | 0 | **219** |
| **sum** | **10** | **15** | **103** | **280** | **439** | **571** | **666** | **719** | **670** | **615** | **519** | **401** | **238** | **105** | **29** | **3** |  |

**Supplementary Table 2**: Descriptive statistics (cognitive performance) of individual tests separately for males and females as well as for all individuals.

| **Test** |  | **T1** | **T2** | **T3** | **T4** | **T5** | **T6** |
| --- | --- | --- | --- | --- | --- | --- | --- |
|  |  | N | N | N | N | N | N |
|  |  | Mean | Mean | Mean | Mean | Mean | Mean |
|  |  | SD | SD | SD | SD | SD | SD |
| Episodic memory  composite | All | 681 | 1324 | 1073 | 864 | 733 | 478 |
|  |  | 35.85 | 35.53 | 37.24 | 37.95 | 38.55 | 38.97 |
|  |  | 9.87 | 11.26 | 10.66 | 10.67 | 10.39 | 10.47 |
|  | Females | 364 | 715 | 579 | 470 | 402 | 258 |
|  |  | 37.06 | 36.52 | 38.68 | 39.84 | 39.85 | 41.16 |
|  |  | 10.12 | 11.67 | 10.71 | 10.73 | 10.65 | 10.21 |
|  | Males | 317 | 609 | 494 | 394 | 331 | 220 |
|  |  | 34.45 | 34.36 | 35.55 | 35.69 | 36.98 | 36.40 |
|  |  | 9.40 | 10.64 | 10.35 | 10.17 | 9.86 | 10.19 |
| Semantic memory  composite | All | 681 | 1337 | 1082 | 871 | 738 | 485 |
|  |  | 21.77 | 21.96 | 22.18 | 23.86 | 23.70 | 24.49 |
|  |  | 8.06 | 8.78 | 8.10 | 8.74 | 8.21 | 8.67 |
|  | Females | 364 | 721 | 584 | 475 | 403 | 262 |
|  |  | 22.20 | 22.90 | 23.02 | 25.20 | 25.02 | 26.39 |
|  |  | 7.90 | 8.85 | 8.15 | 8.87 | 8.34 | 8.26 |
|  | Males | 317 | 616 | 498 | 396 | 335 | 223 |
|  |  | 21.27 | 20.86 | 21.20 | 22.26 | 22.09 | 22.27 |
|  |  | 8.23 | 8.57 | 7.94 | 8.32 | 7.75 | 8.62 |
| Visuospatial | All | 681 | 1331 | 1080 | 863 | 735 | 484 |
| ability |  | 27.98 | 26.98 | 27.23 | 27.66 | 27.94 | 27.34 |
|  |  | 10.45 | 10.70 | 10.77 | 9.86 | 10.03 | 9.87 |
|  | Females | 364 | 718 | 582 | 470 | 403 | 263 |
|  |  | 26.83 | 25.90 | 26.51 | 27.03 | 27.04 | 26.97 |
|  |  | 10.40 | 10.55 | 10.60 | 9.50 | 10.14 | 9.80 |
|  | Males | 317 | 613 | 498 | 393 | 332 | 221 |
|  |  | 29.32 | 28.25 | 28.07 | 28.41 | 29.02 | 27.74 |
|  |  | 10.36 | 10.75 | 10.92 | 10.24 | 9.80 | 9.96 |

SD = standard deviation.

**Supplementary Table 3**: Effect of schizophrenia PRS on cognitive composite cross-sectional (N = 1454), calculated with different p-value thresholds, with corresponding Nagelkerke’s pseudo-R^2^.

|  | **Estimate (95% CI)** | **t-value** | **p-value** | **Nagelkerke’s pseudo-R^2^** |
| --- | --- | --- | --- | --- |
| p ≤ 1 | -0.108 (-0.204 to -0.013) | -2.226 | 0.026* | 0.00350 |
| p < 0.5 | -0.098 (-0.194 to -0.003) | -2.014 | 0.044* | 0.00287 |
| p < 0.4 | -0.102 (-0.197 to -0.006) | -2.089 | 0.037* | 0.00308 |
| p < 0.3 | -0.093 (-0.189 to 0.002) | -1.910 | 0.056 | 0.00258 |
| p < 0.2 | -0.079 (-0.174 to 0.016) | -1.628 | 0.104 | 0.00187 |
| p < 0.1 | -0.077 (-0.172 to 0.019) | -1.578 | 0.115 | 0.00176 |
| p < 0.05 | -0.074 (-0.168 to 0.021) | -1.529 | 0.127 | 0.00165 |
| p < 0.01 | -0.097 (-0.192 to -0.003) | -2.018 | 0.044* | 0.00288 |
| p < 0.001 | -0.071 (-0.165 to 0.023) | -1.478 | 0.140 | 0.00155 |
| p < 0.0001 | -0.045 (-0.138 to 0.050) | -0.910 | 0.363 | 0.00059 |

* = p < 0.05. PRS = polygenic risk score. Linear regression analyses included sex, age, age^2^, sample and the first 5 genetic principal components for genetic ancestry as covariates of no interest, and were based on each individual’s first cognitive test occasion. CI = confidence interval.

**Supplementary Table 4:** Effect of schizophrenia PRS on intercept and slope of cognition composite as well as individual cognitive tests (T1-T6) calculated with different p-value thresholds.

|  |  |  | **Estimate (95% CI)** | **t-value** | **p-value** |
| --- | --- | --- | --- | --- | --- |
| Cognition composite (N = 1454) | p ≤ 1 | PRS | -0.107 (-0.201 to -0.013) | -2.224 | 0.026* |
|  |  | Time*PRS | <0.001 (-0.003 to 0.004) | 0.188 | 0.851 |
|  | p < 0.5 | PRS | -0.096 (-0.200 to -0.002) | -1.990 | 0.047* |
|  |  | Time*PRS | <0.001 (-0.003 to 0.004) | 0.360 | 0.719 |
|  | p < 0.4 | PRS | -0.099 (-0.193 to -0.005) | -2.052 | 0.040* |
|  |  | Time*PRS | <0.001 (-0.003 to 0.004) | 0.374 | 0.709 |
|  | p < 0.3 | PRS | -0.089 (-0.183 to 0.005) | -1.851 | 0.064 |
|  |  | Time*PRS | <0.001 (-0.003 to 0.005) | 0.465 | 0.642 |
|  | p < 0.2 | PRS | -0.076 (-0.170 to 0.018) | -1.584 | 0.113 |
|  |  | Time*PRS | <0.001 (-0.003 to 0.004) | 0.359 | 0.720 |
|  | p < 0.1 | PRS | -0.073 (-0.167 to 0.021) | -1.518 | 0.129 |
|  |  | Time*PRS | <0.001 (-0.003 to 0.004) | 0.115 | 0.908 |
|  | p < 0.05 | PRS | -0.070 (-0.163 to 0.024) | -1.459 | 0.145 |
|  |  | Time*PRS | 0.001 (-0.002 to 0.004) | 0.757 | 0.449 |
|  | p < 0.01 | PRS | -0.095 (-0.188 to -0.002) | -1.986 | 0.047* |
|  |  | Time*PRS | 0.001 (-0.002 to 0.005) | 0.625 | 0.532 |
|  | p < 0.001 | PRS | -0.064 (-0.157 to 0.028) | -1.361 | 0.174 |
|  |  | Time*PRS | <0.001 (-0.003 to 0.005) | 0.550 | 0.582 |
|  | p < 0.0001 | PRS | -0.042 (-0.135 to 0.050) | -0.892 | 0.372 |
|  |  | Time*PRS | 0.002 (-0.001 to 0.005) | 1.194 | 0.232 |
| Episodic memory composite  (N = 1454) | p ≤ 1 | PRS | -0.154 (-0.298 to -0.009) | -2.083 | 0.037* |
|  |  | Time*PRS | -0.002 (-0.009 to 0.005) | -0.526 | 0.599 |
|  | p < 0.5 | PRS | -0.131 (-0.275 to 0.014) | -1.767 | 0.077 |
|  |  | Time*PRS | -0.001 (-0.008 to 0.005) | -0.355 | 0.723 |
|  | p < 0.4 | PRS | -0.136 (-0.280 to 0.009) | -1.838 | 0.066 |
|  |  | Time*PRS | -0.001 (-0.008 to 0.005) | -0.301 | 0.763 |
|  | p < 0.3 | PRS | -0.130 (-0.274 to 0.015) | -1.754 | 0.080 |
|  |  | Time*PRS | -0.001 (-0.008 to 0.006) | -0.231 | 0.817 |
|  | p < 0.2 | PRS | -0.108 (-0.252 to 0.036) | -1.464 | 0.143 |
|  |  | Time*PRS | -0.001 (-0.008 to 0.006) | -0.269 | 0.788 |
|  | p < 0.1 | PRS | -0.028 (-0.257 to 0.030) | -1.548 | 0.122 |
|  |  | Time*PRS | -0.001 (-0.008 to 0.007) | -0.138 | 0.890 |
|  | p < 0.05 | PRS | -0.111 (-0.254 to 0.032) | -1.521 | 0.129 |
|  |  | Time*PRS | 0.004 (-0.003 to 0.011) | 1.053 | 0.292 |
|  | p < 0.01 | PRS | -0.125 (-0.268 to 0.018) | -1.714 | 0.087 |
|  |  | Time*PRS | 0.003 (-0.005 to 0.009) | 0.651 | 0.515 |
|  | p < 0.001 | PRS | -0.097 (-0.239 to 0.045) | -1.335 | 0.182 |
|  |  | Time*PRS | 0.004 (-0.003 to 0.010) | 1.036 | 0.300 |
|  | p < 0.0001 | PRS | -0.057 (-0.199 to 0.085) | -0.782 | 0.434 |
|  |  | Time*PRS | 0.003 (-0.004 to 0.010) | 0.842 | 0.400 |
| Semantic memory composite  (N = 1459) | p ≤ 1 | PRS | -0.122 (-0.226 to -0.017) | -2.276 | 0.023* |
|  |  | Time*PRS | 0.003 (-0.002 to 0.008) | 1.064 | 0.287 |
|  | p < 0.5 | PRS | -0.119 (-0.224 to -0.016) | -2.249 | 0.025* |
|  |  | Time*PRS | 0.003 (-0.002 to 0.008) | 1.156 | 0.248 |
|  | p < 0.4 | PRS | -0.120 (-0.225 to -0.016) | -2.256 | 0.024* |
|  |  | Time*PRS | 0.004 (-0.002 to 0.009) | 1.243 | 0.214 |
|  | p < 0.3 | PRS | -0.118 (-0.223 to -0.015) | -2.228 | 0.026* |
|  |  | Time*PRS | 0.004 (-0.002 to 0.009) | 1.345 | 0.179 |
|  | p < 0.2 | PRS | -0.111 (-0.216 to -0.007) | -2.094 | 0.036* |
|  |  | Time*PRS | 0.004 (-0.002 to 0.009) | 1.287 | 0.198 |
|  | p < 0.1 | PRS | -0.108 (-0.213 to -0.005) | -2.042 | 0.041* |
|  |  | Time*PRS | 0.003 (-0.003 to 0.007) | 0.856 | 0.392 |
|  | p < 0.05 | PRS | -0.090 (-0.191 to 0.013) | -1.700 | 0.089 |
|  |  | Time*PRS | 0.002 (-0.004 to 0.006) | 0.400 | 0.689 |
|  | p < 0.01 | PRS | -0.125 (-0.228 to -0.021) | -2.359 | **0.018*** |
|  |  | Time*PRS | 0.002 (-0.003 to 0.007) | 0.679 | 0.497 |
|  | p < 0.001 | PRS | -0.057 (-0.160 to 0.045) | -1.099 | 0.272 |
|  |  | Time*PRS | <0.001 (-0.004 to 0.006) | 0.306 | 0.760 |
|  | p < 0.0001 | PRS | -0.043 (-0.146 to 0.060) | -0.816 | 0.415 |
|  |  | Time*PRS | <0.001 (-0.004 to 0.006) | 0.314 | 0.753 |
| Visuospatial ability  (N = 1457) | p ≤ 1 | PRS | -0.023 (-0.063 to 0.020) | -1.017 | 0.309 |
|  |  | Time*PRS | -0.001 (-0.002 to 0.001) | -0.481 | 0.631 |
|  | p < 0.5 | PRS | -0.016 (-0.058 to 0.025) | -0.766 | 0.444 |
|  |  | Time*PRS | -0.001 (-0.002 to 0.001) | -0.417 | 0.677 |
|  | p < 0.4 | PRS | -0.017 (-0.059 to 0.024) | -0.824 | 0.410 |
|  |  | Time*PRS | -0.001 (-0.002 to 0.001) | -0.529 | 0.597 |
|  | p < 0.3 | PRS | -0.010 (-0.052 to 0.031) | -0.493 | 0.622 |
|  |  | Time*PRS | -0.001 (-0.002 to 0.001) | -0.550 | 0.582 |
|  | p < 0.2 | PRS | -0.006 (-0.048 to 0.035) | -0.303 | 0.762 |
|  |  | Time*PRS | -0.001 (0.002 to 0.001) | -0.656 | 0.512 |
|  | p < 0.1 | PRS | -0.002 (-0.043 to 0.040) | -0.075 | 0.940 |
|  |  | Time*PRS | -0.001 (-0.002 to 0.001) | -0.709 | 0.478 |
|  | p < 0.05 | PRS | -0.011 (-0.052 to 0.031) | -0.509 | 0.611 |
|  |  | Time*PRS | <0.001 (-0.001 to 0.002) | 0.539 | 0.590 |
|  | p < 0.01 | PRS | -0.018 (-0.059 to 0.024) | -0.835 | 0.404 |
|  |  | Time*PRS | <0.001 (-0.001 to 0.002) | 0.403 | 0.687 |
|  | p < 0.001 | PRS | -0.021 (-0.062 to 0.020) | -1.014 | 0.311 |
|  |  | Time*PRS | <0.001 (-0.001 to 0.002) | 0.278 | 0.781 |
|  | p < 0.0001 | PRS | -0.009 (-0.050 to 0.031) | -0.443 | 0.658 |
|  |  | Time*PRS | 0.002 (-0.0002 to 0.003) | 1.751 | 0.080 |

* = p < 0.05 (uncorrected). Bold = p < 0.02 (corrected for multiple testing). PRS = polygenic risk score. Linear mixed-effect models included sex, age, age^2^ and the first 5 genetic principal components for genetic ancestry as covariates of no interest. CI = confidence interval.

| **Supplementary Table 5**: Effect of schizophrenia PRS on cognition composite as well as individual cognitive tests cross-sectional, calculated with different p-value thresholds, with corresponding interaction with sex. | | | | | |
| --- | --- | --- | --- | --- | --- |
|  |  |  | **Estimate (95% CI)** | **t-value** | **p-value** |
| Cognition composite  (N = 1454) | p ≤ 1 | PRS | -0.110 (-0.205 to -0.015) | -2.260 | 0.024* |
|  |  | Sex*PRS | -0.213 (-0.400 to -0.028) | -2.262 | 0.024* |
|  | p < 0.5 | PRS | -0.099 (-0.194 to -0.003) | -2.030 | 0.043* |
|  |  | Sex*PRS | -0.215 (-0.400 to -0.030) | -2.276 | 0.023* |
|  | p < 0.4 | PRS | -0.103 (-0.198 to -0.007) | -2.114 | 0.035* |
|  |  | Sex*PRS | -0.218 (-0.403 to -0.033) | -2.313 | 0.021* |
|  | p < 0.3 | PRS | -0.094 (-1.190 to 0.002) | -1.930 | 0.054 |
|  |  | Sex*PRS | -0.238 (-0.423 to -0.053) | -2.524 | **0.012*** |
|  | p < 0.2 | PRS | -0.080 (-0.175 to 0.015) | -1.649 | 0.099 |
|  |  | Sex*PRS | -0.224 (-0.409 to -0.038) | -2.368 | **0.018*** |
|  | p < 0.1 | PRS | -0.078 (-0.173 to 0.017) | -1.611 | 0.107 |
|  |  | Sex*PRS | -0.221 (-0.406 to -0.035) | -2.333 | **0.020*** |
|  | p < 0.05 | PRS | -0.075 (-0.170 to 0.020) | -1.557 | 0.120 |
|  |  | Sex*PRS | -0.294 (-0.480 to -0.107) | -3.095 | **0.002*** |
|  | p < 0.01 | PRS | -0.099 (-0.193 to -0.004) | -2.050 | 0.041* |
|  |  | Sex*PRS | -0.274 (-0.460 to -0.089) | -2.899 | **0.004*** |
|  | p < 0.001 | PRS | -0.072 (-0.166 to 0.022) | -1.501 | 0.134 |
|  |  | Sex*PRS | -0.256 (-0.442 to -0.069) | -2.689 | **0.007*** |
|  | p < 0.0001 | PRS | -0.043 (-0.137 to 0.051) | -0.892 | 0.373 |
|  |  | Sex*PRS | -0.195 (-0.380 to -0.009) | -2.059 | 0.040* |
| Episodic  memory  composite  (N = 1454) | p ≤ 1 | PRS | -0.142 (-0.290 to 0.006) | -1.886 | 0.060 |
|  |  | Sex*PRS | -0.269 (-0.555 to 0.018) | -1.840 | 0.066 |
|  | p < 0.5 | PRS | -0.120 (-0.268 to 0.027) | -1.598 | 0.110 |
|  |  | Sex*PRS | -0.251 (-0.537 to 0.036) | -1.717 | 0.086 |
|  | p < 0.4 | PRS | -0.129 (-0.277 to 0.019) | -1.713 | 0.087 |
|  |  | Sex*PRS | -0.255 (-0.541 to 0.031) | -1.746 | 0.081 |
|  | p < 0.3 | PRS | -0.122 (-0.270 to 0.026) | -1.617 | 0.106 |
|  |  | Sex*PRS | -0.302 (-0.588 to -0.016) | -2.070 | 0.039* |
|  | p < 0.2 | PRS | -0.098 (-0.246 to 0.049) | -1.304 | 0.192 |
|  |  | Sex*PRS | -0.307 (-0.594 to -0.021) | -2.103 | 0.036* |
|  | p < 0.1 | PRS | -0.106 (-0.253 to 0.041) | -1.419 | 0.156 |
|  |  | Sex*PRS | -0.304 (-0.590 to -0.017) | -2.080 | 0.038* |
|  | p < 0.05 | PRS | -0.104 (-0.251 to 0.042) | -1.401 | 0.161 |
|  |  | Sex*PRS | -0.302 (-0.590 to -0.014) | -2.055 | 0.040* |
|  | p < 0.01 | PRS | -0.125 (-0.271 to 0.022) | -1.672 | 0.095 |
|  |  | Sex*PRS | -0.322 (-0.610 to -0.035) | -2.198 | 0.028* |
|  | p < 0.001 | PRS | -0.081 (-0.226 to 0.065) | -1.090 | 0.276 |
|  |  | Sex*PRS | -0.189 (-0.478 to 0.100) | -1.282 | 0.200 |
|  | p < 0.0001 | PRS | -0.025 (-0.170 to 0.121) | -0.333 | 0.740 |
|  |  | Sex*PRS | -0.247 (-0.533 to 0.040) | -1.687 | 0.092 |
| Semantic memory composite  (N = 1459) | p ≤ 1 | PRS | -0.131 (-0.239 to -0.023) | -2.375 | **0.018*** |
|  |  | Sex*PRS | -0.160 (-0.369 to 0.050) | -1.492 | 0.136 |
|  | p < 0.5 | PRS | -0.128 (-0.236 to -0.020) | -2.317 | 0.021* |
|  |  | Sex*PRS | -0.167 (-0.377 to 0.043) | -1.562 | 0.119 |
|  | p < 0.4 | PRS | -0.127 (-0.235 to -0.019) | -2.299 | 0.022* |
|  |  | Sex*PRS | -0.169 (-0.379 to 0.040) | -1.585 | 0.113 |
|  | p < 0.3 | PRS | -0.126 (-0.234 to -0.018) | -2.283 | 0.023* |
|  |  | Sex*PRS | -0.187 (-0.397 to 0.023) | -1.749 | 0.080 |
|  | p < 0.2 | PRS | -0.121 (-0.229 to -0.013) | -2.194 | 0.028* |
|  |  | Sex*PRS | -0.187 (-0.397 to 0.023) | -1.744 | 0.081 |
|  | p < 0.1 | PRS | -0.118 (-0.225 to -0.010) | -2.146 | 0.032* |
|  |  | Sex*PRS | -0.201 (-0.411 to 0.009) | -1.873 | 0.061 |
|  | p < 0.05 | PRS | -0.102 (-0.209 to 0.004) | -1.880 | 0.060 |
|  |  | Sex*PRS | -0.235 (-0.446 to -0.025) | -2.192 | 0.029* |
|  | p < 0.01 | PRS | -0.133 (-0.239 to -0.026) | -2.440 | **0.015*** |
|  |  | Sex*PRS | -0.244 (-0.454 to -0.034) | -2.283 | 0.023* |
|  | p < 0.001 | PRS | -0.085 (-0.191 to 0.022) | -1.562 | 0.118 |
|  |  | Sex*PRS | -0.270 (0.481 to -0.060) | -2.522 | **0.012*** |
|  | p < 0.0001 | PRS | -0.058 (-0.164 to 0.049) | -1.061 | 0.289 |
|  |  | Sex*PRS | -0.181 (-0.391 to 0.029) | -1.691 | 0.091 |
| Visuospatial ability  (N = 1457) | p ≤ 1 | PRS | -0.025 (-0.068 to 0.017) | -1.163 | 0.245 |
|  |  | Sex*PRS | -0.079 (-0.162 to 0.004) | -1.860 | 0.063 |
|  | p < 0.5 | PRS | -0.021 (-0.063 to 0.022) | -0.943 | 0.346 |
|  |  | Sex*PRS | -0.082 (-0.165 to 0.001) | -1.931 | 0.054 |
|  | p < 0.4 | PRS | -0.023 (-0.066 to 0.020) | -1.052 | 0.293 |
|  |  | Sex*PRS | -0.083 (-0.166 to 0.0001) | -1.964 | 0.049* |
|  | p < 0.3 | PRS | -0.056 (-0.058 to 0.027) | -0.716 | 0.474 |
|  |  | Sex*PRS | -0.084 (-0.166 to -0.001) | -1.977 | 0.048* |
|  | p < 0.2 | PRS | -0.010 (-0.053 to 0.033) | -0.486 | 0.647 |
|  |  | Sex*PRS | -0.070 (-0.150 to 0.016) | -1.577 | 0.115 |
|  | p < 0.1 | PRS | -0.006 (-0.049 to 0.036) | -0.297 | 0.766 |
|  |  | Sex*PRS | -0.056 (-0.139 to 0.027) | -1.322 | 0.186 |
|  | p < 0.05 | PRS | -0.012 (-0.054 to 0.030) | -0.563 | 0.574 |
|  |  | Sex*PRS | -0.108 (-0.191 to -0.025) | -2.551 | **0.011*** |
|  | p < 0.01 | PRS | -0.018 (-0.060 to 0.025) | -0.818 | 0.414 |
|  |  | Sex*PRS | -0.078 (-0.161 to 0.005) | -1.849 | 0.065 |
|  | p < 0.001 | PRS | -0.025 (-0.067 to 0.017) | -1.182 | 0.237 |
|  |  | Sex*PRS | -0.015 (-0.175 to -0.008) | -2.156 | 0.031* |
|  | p < 0.0001 | PRS | -0.015 (-0.057 to 0.027) | -0.707 | 0.480 |
|  |  | Sex*PRS | -0.057 (-0.140 to 0.026) | -1.344 | 0.179 |

* = p < 0.05 (uncorrected). Bold = p < 0.02 (corrected for multiple testing). PRS = polygenic risk score. Linear regression analyses included sex, age, age^2^, sample and the first 5 genetic principal components for genetic ancestry as covariates of no interest, and were based on each individual’s first cognitive test occasion. CI = confidence interval.

| **Supplementary Table 6:** Effect of schizophrenia polygenic risk score (PRS) (p-value threshold ≤ 1) on intercept and slope of cognition composite as well as individual cognitive tests, while accounting for cognition polygenic score (Cog-PGS) in the model, including all six test occasions, separately for males and females as well as for all individuals. | | | | | |
| --- | --- | --- | --- | --- | --- |
|  |  |  | **Estimate (95% CI)** | **t-value** | **p-value** |
| Cognition composite | PRS | All  (N =  1454) | -0.076 (-0.168 to -0.015) | -1.626 | 0.1041 |
|  | Time*PRS |  | 0.001 (-0.003 to 0.004) | 0.252 | 0.8011 |
|  | Cog-PGS |  | 0.406 (0.316 to 0.496) | 8.819 | **<2e-16*** |
|  | Time*Cog-PGS |  | 0.001 (-0.003 to 0.005) | 0.580 | 0.5619 |
|  | PRS | Females  (N =  778) | 0.183 (-0.108 to 0.145) | 0.282 | 0.7776 |
|  | Time*PRS |  | 0.001 (-0.004 to 0.006) | 0.336 | 0.7367 |
|  | Cog-PGS |  | 0.443 (0.320 to 0.566) | 7.014 | **4.58e-12*** |
|  | Time*Cog-PGS |  | 0.001 (-0.005 to 0.005) | 0.067 | 0.9463 |
|  | PRS | Males  (N =  676) | -0.180 (-0.312 to -0.047) | -2.632 | **0.0087*** |
|  | Time*PRS |  | 0.001 (-0.005 to 0.006) | 0.124 | 0.9013 |
|  | Cog-PGS |  | 0.360 (0.228 to 0.492) | 5.325 | **1.33e-07*** |
|  | Time*Cog-PGS |  | 0.002 (-0.003 to 0.007) | 0.780 | 0.4355 |
| Episodic memory composite | PRS | All  (N =  1450) | -0.119 (-0.261 to 0.024) | -1.629 | 0.1036 |
|  | Time*PRS |  | -0.002 (-0.009 to 0.005) | -0.487 | 0.6259 |
|  | Cog-PGS |  | 0.461 (0.322 to 0.601) | 6.459 | **1.35e-10*** |
|  | Time*Cog-PGS |  | 0.001 (-0.006 to 0.008) | 0.270 | 0.7869 |
|  | PRS | Females  (N =  777) | 0.002 (-0.195 to 0.199) | 0.022 | 0.9824 |
|  | Time*PRS |  | -0.002 (-0.012 to 0.008) | -0.385 | 0.7000 |
|  | Cog-PGS |  | 0.485 (0.294 to 0.677) | 4.933 | **9.52e-07*** |
|  | Time*Cog-PGS |  | 0.002 (-0.008 to 0.011) | 0.361 | 0.7183 |
|  | PRS | Males  (N =  673) | -0.248 (-0.452 to -0.043) | -2.361 | **0.0185*** |
|  | Time*PRS |  | -0.001 (-0.011 to 0.009) | -0.232 | 0.8164 |
|  | Cog-PGS |  | 0.437 (0.235 to 0.640) | 4.200 | **2.96e-05*** |
|  | Time*Cog-PGS |  | -0.001 (-0.010 to 0.010) | -0.066 | 0.9477 |
| Semantic memory composite | PRS | All  (N =  1458) | -0.092 (-0.194 to 0.011) | -1.740 | 0.0820 |
|  | Time*PRS |  | 0.003 (-0.002 to 0.008) | 1.051 | 0.2932 |
|  | Cog-PGS |  | 0.381 (0.281 to 0.482) | 7.408 | **1.92e-13*** |
|  | Time*Cog-PGS |  | -0.001 (-0.006 to 0.005) | -0.182 | 0.8552 |
|  | PRS | Females  (N =  781) | -0.037 (-0.176 to 0.103) | -0.516 | 0.6056 |
|  | Time*PRS |  | 0.004 (-0.003 to 0.011) | 1.080 | 0.2804 |
|  | Cog-PGS |  | 0.444 (0.309 to 0.580) | 6.397 | **2.41e-10*** |
|  | Time*Cog-PGS |  | -0.002 (-0.009 to 0.005) | -0.569 | 0.5691 |
|  | PRS | Males  (N =  677) | -0.154 (-0.306 to -0.003) | -1.986 | 0.0474* |
|  | Time*PRS |  | 0.002 (-0.006 to 0.009) | 0.470 | 0.6384 |
|  | Cog-PGS |  | 0.302 (0.152 to 0.453) | 3.921 | **9.53e-05*** |
|  | Time*Cog-PGS |  | 0.002 (-0.006 to 0.009) | 0.458 | 0.6471 |
| Visuospatial ability | PRS | All  (N =  1456) | -0.011 (-0.053 to 0.030) | -0.538 | 0.5906 |
|  | Time*PRS |  | -0.001 (-0.002 to 0.001) | -0.410 | 0.6818 |
|  | Cog-PGS |  | 1.339 (0.093 to 0.174) | 6.473 | **1.25e-10*** |
|  | Time*Cog-PGS |  | 0.001 (-0.001 to 0.002) | 0.684 | 0.4943 |
|  | PRS | Females  (N =  780) | 0.032 (-0.024 to 0.089) | 1.116 | 0.2647 |
|  | Time*PRS |  | -0.001 (-0.003 to 0.003) | -0.394 | 0.6934 |
|  | Cog-PGS |  | 0.146 (0.909 to 0.200) | 5.189 | **2.60e-07*** |
|  | Time*Cog-PGS |  | 0.001 (-0.002 to 0.003) | 0.304 | 0.7615 |
|  | PRS | Males  (N =  676) | -0.058 (-0.119 to 0.003) | -1.863 | 0.0629 |
|  | Time*PRS |  | -0.001 (-0.003 to 0.002) | -0.091 | 0.9273 |
|  | Cog-PGS |  | 0.120 (0.060 to 0.181) | 3.888 | **0.0001*** |
|  | Time*Cog-PGS |  | 0.001 (-0.002 to 0.003) | 0.634 | 0.5260 |

* = p < 0.05 (uncorrected). Bold = p < 0.02 (corrected for multiple testing). Linear mixed-effect models included sex (when applicable), age, age^2^, the COG-PGS for cognitive performance, and the first 5 genetic principal components for genetic ancestry as covariates of no interest. CI = confidence interval.

| **Supplementary Table 7:** Effect of schizophrenia polygenic risk score (PRS) (p-value threshold ≤ 1) on intercept and slope of cognition composite as well as individual cognitive tests, while accounting for years of education (Educ) in the model, including all six test occasions, separately for males and females as well as for all individuals. | | | | | |
| --- | --- | --- | --- | --- | --- |
|  |  |  | **Estimate (95% CI)** | **t-value** | **p-value** |
| Cognition composite | PRS | All  (N =  1454) | -0.114 (-0.202 to -0.025) | -2.512 | **0.0121*** |
|  | Time*PRS |  | -0.001 (-0.004 to 0.003) | -0.154 | 0.8775 |
|  | Educ |  | 0.671 (0.573 to 0.769) | 13.44 | **<2e-16*** |
|  | Time*Educ |  | -0.007 (-0.011 to -0.003) | -3.095 | **0.0020*** |
|  | PRS | Females  (N =  778) | -0.022 (-0.146 to 0.032) | -0.332 | 0.7401 |
|  | Time*PRS |  | 0.001 (-0.005 to 0.002) | 0.094 | 0.9249 |
|  | Educ |  | 0.568 (0.436 to 0.696) | 8.495 | **<2e-16*** |
|  | Time*Educ |  | -0.004 (-0.010 to 0.002) | -1.380 | 0.1678 |
|  | PRS | Males  (N =  676) | -0.199 (-0.322 to -0.075) | -3.125 | **0.0019*** |
|  | Time*PRS |  | -0.001 (-0.006 to 0.004) | -0.308 | 0.7582 |
|  | Educ |  | 0.835 (0.686 to 0.985) | 10.97 | **<2e-16*** |
|  | Time*Educ |  | -0.107 (-0.017 to -0.004) | -3.138 | **0.0017*** |
| Episodic memory composite | PRS | All  (N =  1450) | -0.163 (-0.301 to -0.025) | -2.312 | 0.0209* |
|  | Time*PRS |  | -0.002 (-0.010 to 0.005) | -0.669 | 0.5036 |
|  | Educ |  | 0.903 (0.746 to 1.062) | 11.175 | **<2e-16*** |
|  | Time*Educ |  | -0.003 (-0.010 to 0.005) | -0.796 | 0.4262 |
|  | PRS | Females  (N =  777) | -0.048 (-0.240 to 0.143) | -0.492 | 0.6231 |
|  | Time*PRS |  | -0.003 (-0.013 to 0.007) | 0.288 | 0.5502 |
|  | Educ |  | 0.831 (0.622 to 1.042) | 7.736 | **2.03e-14*** |
|  | Time*Educ |  | 0.002 (-0.010 to 0.014) | 0.288 | 0.7734 |
|  | PRS | Males  (N =  673) | -0.273 (-0.469 to -0.077) | -2.708 | **0.0069*** |
|  | Time*PRS |  | -0.002 (-0.012 to 0.008) | -0.347 | 0.7287 |
|  | Educ |  | 1.027 (0.785 to 1.270) | 8.279 | **3.64e-16*** |
|  | Time*Educ |  | -0.010 (-0.022 to 0.002) | -1.583 | 0.1136 |
| Semantic memory composite | PRS | All  (N =  1458) | -0.127 (-0.228 to -0.027) | -2.473 | **0.0135*** |
|  | Time*PRS |  | 0.002 (-0.003 to 0.008) | 0.932 | 0.3684 |
|  | Educ |  | 0.606 (0.491 to 0.723) | 10.244 | **<2e-16*** |
|  | Time*Educ |  | -0.003 (-0.009 to 0.003) | -0.900 | 0.3684 |
|  | PRS | Females  (N =  781) | -0.074 (-0.213 to 0.065) | -1.039 | 0.2993 |
|  | Time*PRS |  | 0.004 (-0.004 to 0.011) | 0.975 | 0.3297 |
|  | Educ |  | 0.468 (0.316 to 0.621) | 5.990 | **2.68e-09*** |
|  | Time*Educ |  | -0.001 (-0.008 to 0.008) | -0.069 | 0.9450 |
|  | PRS | Males  (N =  677) | -0.169 (-0.313 to -0.025) | -2.280 | 0.0229* |
|  | Time*PRS |  | 0.001 (-0.006 to 0.003) | 0.309 | 0.7574 |
|  | Educ |  | 0.813 (0.635 to 0.992) | 8.880 | **<2e-16*** |
|  | Time*Educ |  | -0.006 (-0.016 to 0.003) | -1.328 | 0.1843 |
| Visuospatial ability | PRS | All  (N =  1456) | -0.023 (-0.064 to 0.017) | -1.114 | 0.2653 |
|  | Time*PRS |  | -0.001 (-0.003 to 0.001) | -0.780 | 0.4354 |
|  | Educ |  | 0.226 (0.181 to 0.272) | 9.769 | **<2e-16*** |
|  | Time*Educ |  | -0.005 (-0.007 to -0.003) | -4.322 | **1.58e-05*** |
|  | PRS | Females  (N =  780) | 0.017 (-0.039 to 0.074) | 0.608 | 0.5433 |
|  | Time*PRS |  | -0.001 (-0.007 to 0.002) | -0.571 | 0.5682 |
|  | Educ |  | 0.209 (0.149 to 0.268) | 6.827 | **1.31e-11*** |
|  | Time*Educ |  | -0.004 (-0.007 to 0.001) | -2.608 | **0.0092*** |
|  | PRS | Males  (N =  676) | -0.064 (-0.123 to -0.005) | -2.102 | 0.0358* |
|  | Time*PRS |  | -0.001 (-0.003 to 0.002) | -0.518 | 0.9273 |
|  | Educ |  | 0.254 (0.183 to 0.325) | 6.991 | **4.75e-12*** |
|  | Time*Educ |  | -0.006 (-0.009 to -0.003) | -3.525 | **0.0004*** |

* = p < 0.05 (uncorrected). Bold = p < 0.02 (corrected for multiple testing). Linear mixed-effect models included sex (when applicable), age, age^2^, year of education, and the first 5 genetic principal components for genetic ancestry as covariates of no interest. CI = confidence interval.
